# Pharmacological inhibition of the DNA damage checkpoint prevents radiation-induced oocyte death

**DOI:** 10.1101/132779

**Authors:** Vera D. Rinaldi, Kristin Hsieh, Robert Munroe, Ewelina Bolcun-Filas, John C. Schimenti

**Affiliations:** Dept. of Biomedical Sciences, Cornell University, College of Veterinary Medicine, Ithaca, New York; Dept. of Molecular Biology and Genetics, Cornell University, Ithaca, New York

**Keywords:** fertility, primordial follicles, premature ovarian failure, oncofertility

## Abstract

Ovarian function is directly correlated with survival of the primordial follicle reserve. Women diagnosed with cancer have a primary imperative of treating the cancer, but since the resting oocytes are hypersensitive to the DNA-damaging modalities of certain chemo- and radiotherapeutic regimens, such patients face the collateral outcome of premature loss of fertility and ovarian endocrine function. Current options for fertility preservation primarily include collection and cryopreservation of oocytes or *in vitro* fertilized oocytes, but this necessitates a delay in cancer treatment and additional assisted reproductive technology (ART) procedures. Here, we evaluated the potential of pharmacological preservation of ovarian function by inhibiting a key element of the oocyte DNA damage checkpoint response, checkpoint kinase 2 (CHK2; CHEK2). Whereas non-lethal doses of ionizing radiation (IR) eradicate immature oocytes in wild type mice, irradiated *Chk2*^*-/-*^ mice retain their oocytes and thus, fertility. Using an ovarian culture system, we show that transient administration of the CHK2 inhibitor 2-(4-(4-Chlorophenoxy)phenyl)-1H-benzimidazole-5-carboxamide-hydrate (“CHK2iII”) blocked activation of the CHK2 targets TRP53 and TRP63 in response to sterilizing doses of IR, and preserved oocyte viability. After transfer into sterilized host females, these ovaries proved functional and readily yielded normal offspring. These results provide experimental evidence that chemical inhibition of CHK2 is a potentially valid treatment for preserving fertility and ovarian endocrine function of women exposed to DNA-damaging cancer therapies such as IR.

**Summary sentence:** Oocytes, which are highly sensitive to DNA damage caused by certain cancer treatments, can be protected from radiation-induced death by an inhibitor of the checkpoint protein CHK2.

## INTRODUCTION

It is of paramount importance that organisms minimize the transmission of deleterious mutations to their offspring. Accordingly, sensitive mechanisms have evolved to eliminate germ cells that have sustained certain threshold amounts of DNA damage (HEYER *et al.* 2000; SUH *et al.* 2006; BOLCUN-FILAS *et al.* 2014; PACHECO *et al.* 2015). Under normal circumstances, this is desirable. However, because women are born with a finite number of oocytes, environmental factors that cause DNA damage to oocytes can result in primary ovarian insufficiency (POI), sterility, and ovarian failure. This is a crucial issue for cancer patients undergoing certain types of chemotherapy or radiation therapy (WOODRUFF 2007). For example, POI occurs in nearly 40% of all female breast cancer survivors (OKTAY *et al.* 2006). The resulting premature ovarian failure has major impact in a women’s life, both physiologically and emotionally. As the life expectancy of cancer survivors increases, so does the need to address the adverse outcomes to fertility. Therefore, the ability to inhibit oocyte death and preserve fertility, in both pre-pubertal cancer patients and premenopausal women, would have a major impact on survivors’ lives.

At present, cancer patients have few options regarding fertility preservation (“oncofertility”) before treatment, and most involve invasive surgical procedures such as extraction of oocytes or ovarian tissue for cryopreservation, or IVF followed by embryo cryopreservation (REDIG *et al.* 2011; SALAMA AND MALLMANN 2015; KIM *et al.* 2016). Not only are these invasive, but also they necessitate a delay in cancer treatment. An alternative is to co-administer drugs that protect oocytes from chemotherapy at the time of treatment. For example, based on the knowledge that inactivation of the “TA” isoform of the DNA damage checkpoint gene *Trp63* (*TP63* in humans, also known as *p63;* the TA isoform of the protein will be referred to as TAp63) prevents radiation-induced oocyte loss in mice (SUH *et al.* 2006), it was reported (GONFLONI *et al.* 2009), but later challenged (KERR *et al.* 2012) and counter-argued (MAIANI *et al.* 2012), that the tyrosine kinase inhibitor imatinib (Gleevec) is effective in protecting oocytes. Even if imatinib proves to have such activity, it is a relatively promiscuous kinase inhibitor that blocks, among other targets, the receptor tyrosine kinase KIT that functions in germline stem cells (LEE AND WANG 2009).

We previously reported that mouse CHK2 is a key component of the meiotic DNA damage checkpoint, and that deletion of *Chk2* prevented irradiation-induced killing of postnatal oocytes (BOLCUN-FILAS *et al.* 2014). We also showed that the CHK2 kinase phosphorylates both p53 (formally TRP53; TP53 in humans) and TAp63 in oocytes to activate these proteins (and stabilize TRP53). Therefore, the deletion of *Chk2* effectively impairs the activation of these two downstream effectors, which are both needed to trigger efficient oocyte elimination (BOLCUN-FILAS *et al.* 2014). More importantly, damaged oocytes that survived in the absence of CHK2 produced healthy pups suggesting that the inflicted DNA damage was repaired (BOLCUN-FILAS *et al.* 2014). The resistance of *Chk2*^*-/-*^ oocytes to otherwise lethal levels of ionizing radiation (IR) prompted us to explore whether chemical inhibition of CHK2 would be effective at preventing radiation-induced oocyte death, and thus constitute a potential option for preserving ovarian function in women undergoing cancer therapy. Here we show that transient chemical inhibition of CHK2 prevents follicle loss and allows for the production of healthy offspring.

### Results and Discussion

Irradiation of ovaries induces CHK2-dependent phosphorylation of TAp63 in oocytes, and this phosphorylation is essential for triggering their death (SUH *et al.* 2006; BOLCUN-FILAS *et al.* 2014). CHK2 is a key component of the DNA damage response pathway that responds primarily to DNA double strand breaks (DSBs), lying downstream of the apical kinase ATM (ataxia telangiectasia mutated). Members of the ATM>CHK2>p53/p63 pathway have been implicated as potential anti-cancer drug targets sensitizing cancer cells to genotoxic therapies, and chemical inhibitors have been developed against CHK2 (GARRETT AND COLLINS 2011), which, when deleted in mice, causes only minor phenotypic consequences (TAKAI *et al.* 2002). We therefore tested whether a well characterized CHK2 inhibitor 2-(4-(4-Chlorophenoxy)phenyl)-1H-benzimidazole-5-carboxamide-hydrate (designated “Chk2 inhibitor II” by the manufacturer, referred to here as “CHK2iII”) could mimic the oocyte-protective effect of genetic *Chk2* deletion, and if it could do so in a non-toxic manner.

We employed an organ culture paradigm to control dosages, penetration and timing of drug delivery to the ovary. Using dose ranges based upon published data (ARIENTI *et al.* 2005) and the manufacturer’s recommendations, we first tested the ability of CHK2iII to block phosphorylation of TAp63 in irradiated ovaries, and to block the stabilization of p53, which is normally rapidly degraded in cells unless stabilized by DNA-damage-induced phosphorylation by proteins including CHK2 (CHEHAB *et al.* 2000; HIRAO *et al.* 2000). We used ovaries from 5 days postpartum (dpp) females to ensure that oocytes were in the dictyate arrest stage of meiosis, residing within primordial follicles. Explanted ovaries were cultured for two hours in the presence of 0, 10 or 20μM CHK2iII, then subjected (or not) to 3 Gy of IR, a level that not only kills oocytes, but also causes extensive TRP53 stabilization and TAp63 phosphorylation (SUH *et al.* 2006; BOLCUN-FILAS *et al.* 2014). Ovaries were harvested three hours later for protein extraction and western blot analysis. In non-irradiated ovaries, TAp63 remained unphosphorylated and TRP53 was undetectable (Fig. 1). Irradiation in the absence of inhibitor led to robust TRP53 stabilization, and all TAp63 was shifted to a higher mobility, which is known to be due to phosphorylation (SUH *et al.* 2006; LIVERA *et al.* 2008). Addition of 10μM and 20μM CHK2iII led to partial and complete inhibition of TAp63 phosphorylation, respectively, and also progressively decreased TRP53 levels (Fig. 1). This confirms that CHK2iII treatment rapidly acts to prevent activation of two pro-apoptotic factors in the ovary.

**Figure 1.**
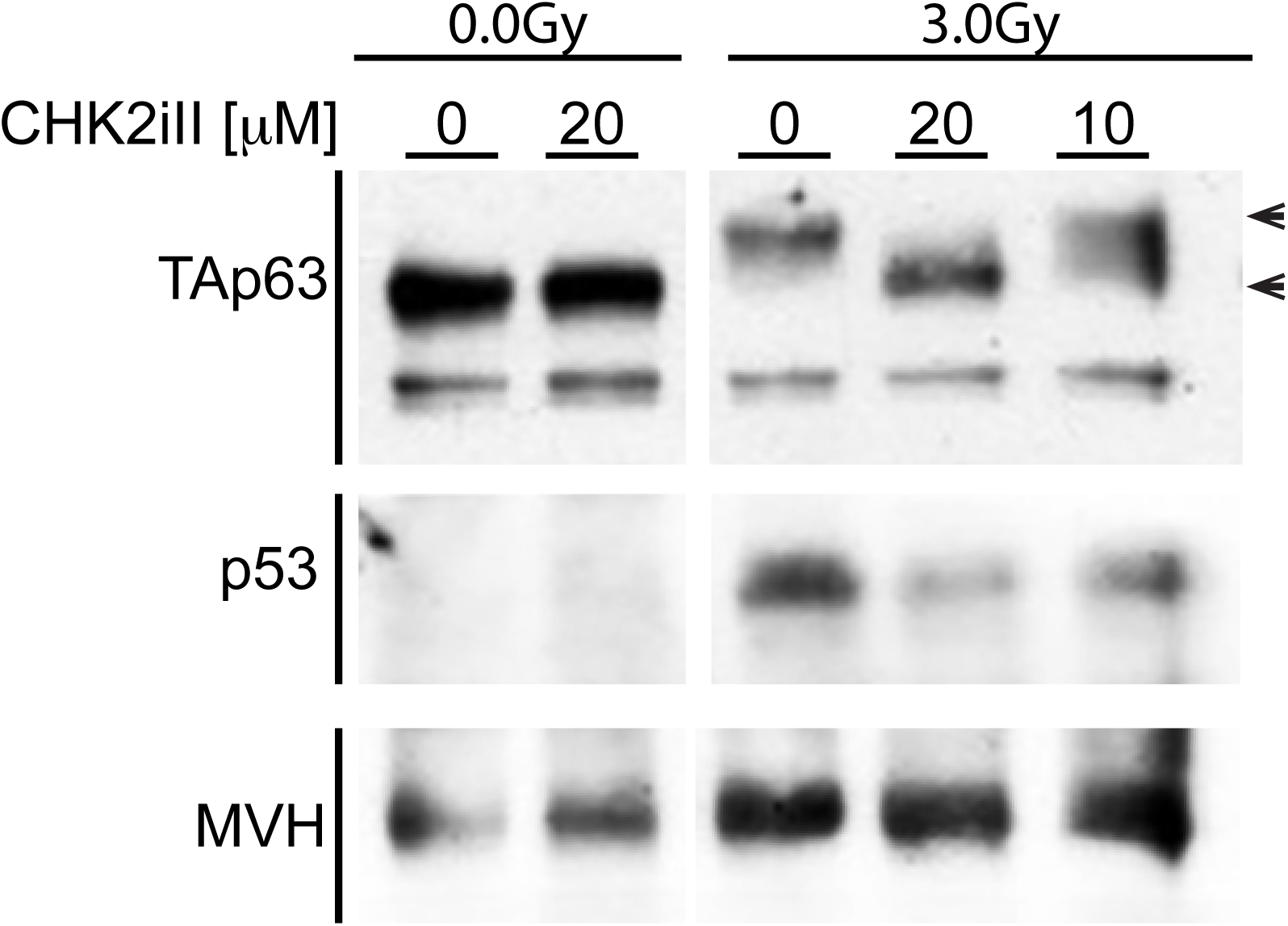
Inhibition of radiation-induced phosphorylation of p53 and TAp63 by CHK2iII. Western blot analysis of protein extracted from five dpp ovaries. Ovaries were incubated with the indicated concentrations of CHK2 inhibitor II (see Methods), and exposed or not to 3 Gy of γ-radiation. The immunoblot membrane was cut into two parts - one containing proteins >60 kDa, and the other <60kDa – and probed with anti-p63 and anti-p53, respectively. The >60 kDa portion was stripped and re-probed for the germ cell marker MVH. Arrowheads indicated the expected molecular weight of the phosphorylated form of TAp63 (upper), and non-phosphorylated TAp63 (lower).

Next, we tested whether CHK2iII could permanently protect oocytes from a lower dose of radiation (0.4 Gy) that normally kills all oocytes within 2 days (SUH *et al.* 2006; BOLCUN-FILAS *et al.* 2014) but is far below levels (>5 Gy) lethal to whole animals. Ovaries were cultured in media supplemented with 0, 5, 10, or 20 μM of inhibitor for 2 hours before irradiation. Following IR exposure, ovaries were cultured 2 more days with media changes as delineated in Fig. 2a, after which the drug was removed from the medium. This protocol of media changes with drug replenishment was optimized for oocyte survival. Seven days after irradiation (and 5 days after removal of CHK2iII, oocyte survival was assessed by co-immunolabeling of histological sections (Fig. S1) and of whole mounts, with the cytoplasmic germ cell marker MVH and oocyte nuclear marker p63 (Figs. 2b, S2a, and Supplemental movies M1, M2, M3). Under these conditions the inhibitor was well tolerated, and oocyte survival in unirradiated ovaries was not compromised (Fig. S2b and Supplemental movies M4, M5).

**Figure 2.**
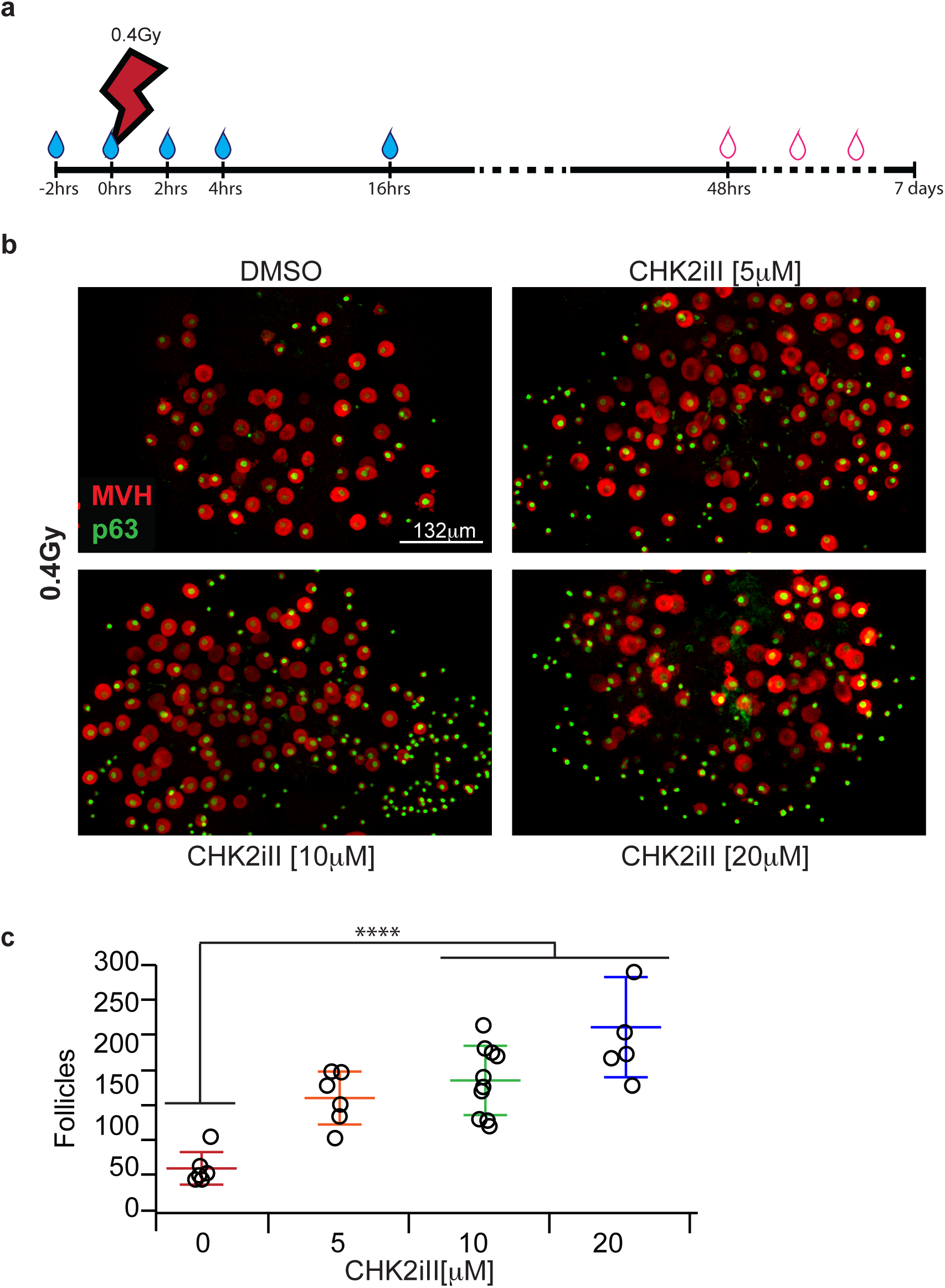
Concentration-dependent protection of irradiated oocytes by CHK2iII. (**a**) Schematic of CHK2iII treatment regimen, beginning with placing explanted 5 dpp ovaries into culture. Blue droplets denote times at which fresh media containing CHK2iII was added/replaced. Red outlined droplets indicate changes with drug-free media. (**b**) Maximum intensity projections of immunostained ovary whole ovaries. For 3D visualization, see Supplementary movies M1, M2 and M3). The ovaries were cultured according to the timeline in “**a”** in the presence of the indicated concentrations of CHK2iII. DMSO corresponds to diluent control. MVH is a cytoplasmic germ cell protein, and p63 labels oocyte nuclei. Note that growing follicles (oocytes with larger MVH-stained cytoplasm) are relatively refractory to IR. (**c**) Quantification of follicles. Data points represent total follicle counts derived from one ovary. Horizontal hashes represent mean and standard deviation. Colors correspond to the different concentrations of inhibitors. Asterisks indicate p-value ≤ 0.0001 (Tukey HSD).

Remarkably, though 10μM CHK2iII only partially inhibited TAp63 phosphorylation induced by 3Gy of IR (Fig. 1), it dramatically improved oocyte survival in ovaries exposed to 0.4Gy of IR, a level sufficient to trigger TAp63 phosphorylation and eliminate nearly all primordial follicles in ovaries (Fig 2b,c; S3) (SUH *et al.* 2006; BOLCUN-FILAS *et al.* 2014). A small but significant protective effect was also observed with 5μM CHK2iII (p=004, Fig. 2b,c).

To assess if oocytes protected by CHK2iII from otherwise lethal levels of irradiation were capable of ovulation, fertilization, and subsequent embryonic development, we performed intrabursal transfers of irradiated ovaries (0.4 Gy) into histocompatible (strain C3H) agouti females. These recipient females were first sterilized at 7 dpp by exposure to 0.5Gy of IR. Once females were eight weeks old, sterility was verified by housing them with fertile males for at least eight weeks. The IR-induced oocyte death led to premature ovarian failure, yielding sufficient intrabursal space without the need to physically remove the vestigial ovaries before ovary transfer. These recipients were 16 weeks of age at the time of surgery. Three cultured ovaries, derived from black female animals also of strain C3H (see Methods for additional information), were placed into each bursa. Fig. 3a summarizes the experimental timeline. A total of 8 successful embryo transfer surgeries were performed. Three females received mock-treated (cultured in media containing 1% DMSO alone) irradiated ovaries (0.4 Gy), and five females received irradiated ovaries (also 0.4 Gy) treated with 10μM CHK2iII in media containing 1% DMSO.

**Figure 3.**
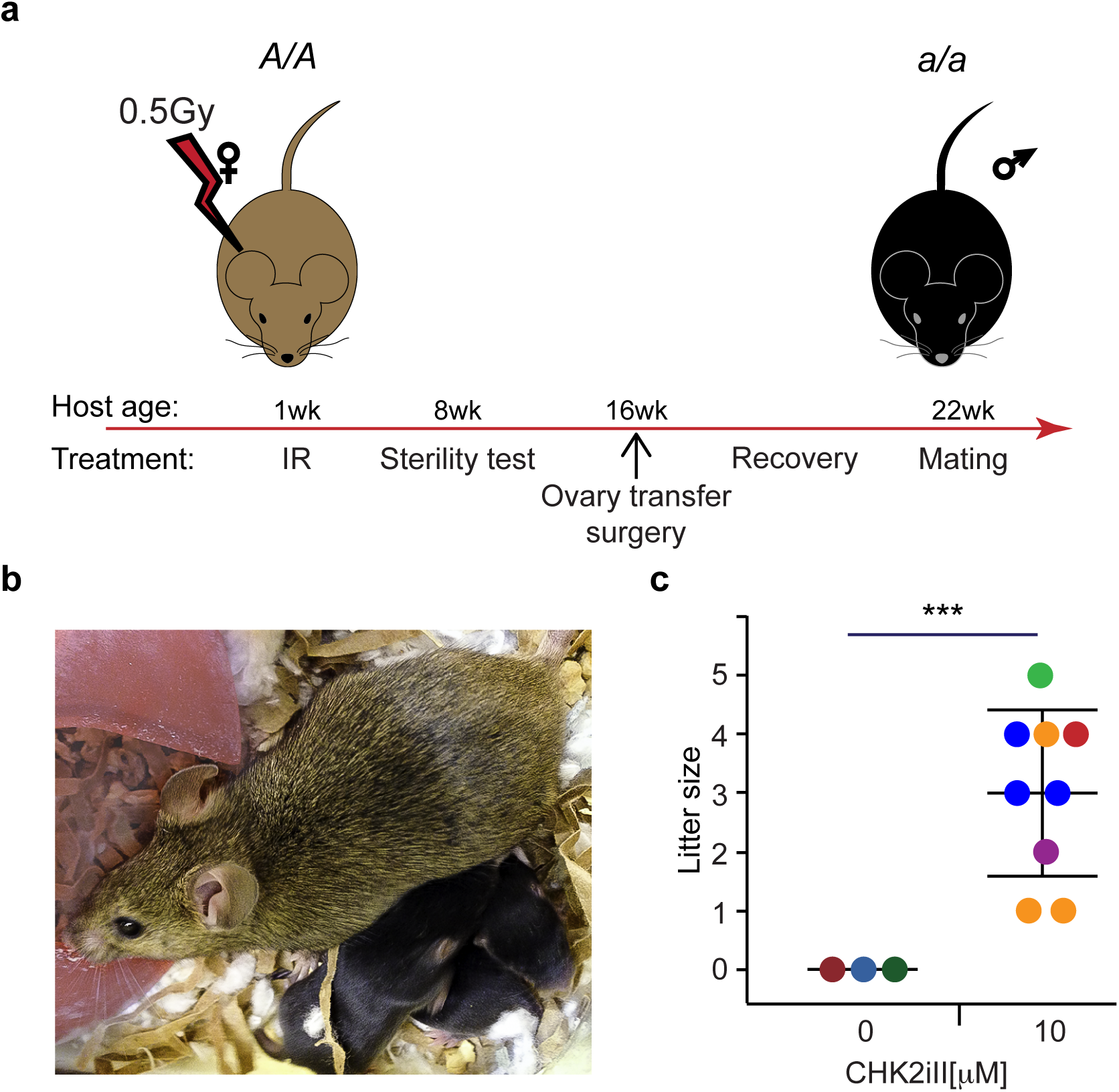
CHK2iII-rescued ovaries are fertile. (**a**) Experimental timeline. Agouti females (*A/A*) were sterilized with 0.5Gy of IR at one week of age. The transplanted ovaries were from black donor females (*a/a*) treated as outlined in Fig. 2a. (**b**) Agouti host females gave birth to black offspring (*a/a*) exclusively; thus, the ovulated eggs produced were from the donor ovaries. (**c**) Litter sizes of mock-treated and treated ovaries. Each circle represents a litter and the circle’s color represents the female that generated that litter. The combined average litter size produced by all host females was 3.

Once the transferred donor ovary reached 8 weeks of age (with respect to the time at which it was explanted), the recipient females were mated to proven fertile C3H black (*a/a*) males for three months, and monitored for litters and the coat colors of offspring. Only females that received CHK2iII-treated ovaries delivered progeny, all of which were black, confirming that they were produced from fertilization of oocytes ovulated from donor ovaries (Fig. 3b and c). All offspring had no visible abnormalities that would suggest inheritance of gross chromosomal abnormalities (BOLCUN-FILAS *et al.* 2014). The viability of these animals indicated that even though oocytes have sensitive checkpoint mechanisms rendering them vulnerable to low levels of DNA damage, they are capable of repairing damaged DNA in a manner compatible with normal embryogenesis.

Our results provide proof-in-principle for the strategy of targeting the CHK2-dependent DNA damage checkpoint pathway for preventing loss of ovarian reserve - and thus ovarian failure in prepubertal or premenopausal cancer patients undergoing therapies that are toxic to oocytes. Importantly, checkpoint inhibitors have already been explored as potential anticancer therapies, thus substantial information is already available on members of this drug class (ANTONI *et al.* 2007; GARRETT AND COLLINS 2011). Nevertheless, it remains to be seen whether systemic administration of CHK2iII, or other CHK2 inhibitors, can achieve similar oocyte-protective efficacy against IR- or drug-induced DSBs *in vivo*. Additionally, it will be important to conduct more thorough studies of potential genetic risks associated oocytes rescued from DNA damage-induced death by checkpoint inhibition.

## Acknowledgements

We thank C. Abratte, Jordana Bloom, Johanna DelaCruz and Rebecca Williams for technical assistance. This work was supported by NIH grants S10OD018516 (to Cornell’s Imaging facility) and R01GM45415 to JCS.

## MATERIALS AND METHODS

### Mice

Mice were obtained from The Jackson Laboratory, strains C3HeB/FeJ, stock # 000658 (agouti mice, homozygous dominant for coat color (*A/A*)) and C3FeLe.B6-a/J, stock # 000198 (black mice, recessive for coat color, (*a/a*)). Cornell’s Animal Care and Use Committee approved all animal usage, under protocol 2004-0038 to JCS.

### Organ Culture

Ovaries were cultured using an adaptation of a published method (LIVERA *et al.* 2008). Briefly, ovaries were collected from five day postpartum (dpp) C3FeLe.B6-a/J mice, and, following removal from the bursa, placed into cell culture inserts (Millicell; pore size: 8μm; diameter: 12mm) pre-soaked in ovary culture media: MEM supplemented with 10% FBS, 25mM HEPES pH=7.0, 100 units/ml penicillin, 100 μg/ml streptomycin, 0.25 μg/ml Fungizone, 1%DMSO, and CHK2 inhibitor. The inserts were placed into 24 well plates (Model MD24 ThermoFisher) with carriers for the inserts. Sufficient media was added to keep organs moist, but not completely submerged. Organs were incubated at 37°C, 5% CO_2_ and atmospheric O_2_.

### Drug Treatment

CHK2iII (CalBiochem 220486) was prepared as 1mM and 2mM stock solutions in DMSO and kept frozen at −20°C. Media containing the desired concentration of inhibitor was prepared shortly before use, assuring that the DMSO concentration was constant (1%DMSO) throughout the different conditions. Explanted ovaries were pre-incubated for 2 hours in warm (37°C) media containing the desired concentration of inhibitor or 1% DMSO alone before being subjected to ionizing radiation in a ^137^cesium irradiator with a rotating turntable. Figure 2a presents the media change regimen, with the first replacement being immediately after irradiation. The ovaries were cultured for either 3 hours before being processed for western blot analysis (to detect DNA damage responses), or 7 days followed by either fixation and immunostaining (to quantify oocyte survival), or ovary transplant surgery into sterile agouti females.

### Western Blot Analyses and Antibodies

Ovary protein lysates, immunoblotting, probing and detection were conducted as described (BOLCUN-FILAS *et al.* 2014). Primary antibodies and dilutions used were: mouse anti-p63 (1:500, 4A4, Novus Biologicals); rabbit anti-p53 (1:300, Cell Signaling #9282); mouse anti-β-actin (1:5000, Sigma) and rabbit anti-MVH (1:1000, Abcam). Secondary antibodies used were: Immuno Pure goat anti-mouse IgG(H+L) peroxidase conjugate (1:5000, ThermoFisher) and goat anti-rabbit IgG HRP-linked antibody (1:5000, Cell Signaling).

### Immunofluorescence

Cultured ovaries were fixed in 4% paraformaldehyde/PBS at 4°C overnight, and then washed and stored in 70% ethanol. Ovaries were either embedded in paraffin and sectioned at 5μm for immunostaining or subjected to whole mount immunostaining and clearing. For the standard immunofluorescence, slides were deparaffinized and rehydrated prior to antigen retrieval using sodium citrate buffer. Slides were blocked with 5% goat serum (PBS/Tween 20), incubated at 4°C overnight with aforementioned primary antibodies (anti-p63 @ 1:500; anti-MVH @ 1:1000), and subsequently incubated with Alexa Fluor® secondary antibodies for one hour and Hoechst dye for 5 minutes. Slides were mounted with ProLong Anti-fade (Thermo-Fisher) and imaged.

### Ovary transfer surgery

Agouti females were radio-sterilized with 0.5Gys of radiation at one week of age. Once eight weeks old, females were housed with males known to be fertile. After eight weeks, and two days prior to ovary transfer surgery, males were removed from female cage. During surgery, three ovaries either treated with 10mM of inhibitor or with 1%DMSO, as described previously in the drug treatment section, were placed in the intrabursal space of each ovary. Females were allowed a recovery period of six weeks. The recovery period was determined as being the amount of time needed for the implanted ovaries (relative age of 12 days old) to reach reproductive age if never remover from their physiological environment. After the recovery period females were house with black males. At 40 weeks of age, upon euthanasia, dissection was performed for visual confirmation of the presence of ovaries for both mock treated and treated.

### Whole organ immunofluorescence

Ovaries cultured for seven days were fixed in freshly prepared 4%paraformaldehyde (PFA) /PBS at 4°C overnight (ON). Afterwards, tissues were washed and stored in 70% ethanol at 4°C until further processing.

Fixed ovaries were washed and left to equilibrate for a minimum of four hours in PBS before initializing whole-mount immunostaining protocol. In order to facilitate handling and tissue integrity, ovaries were kept in the culture insert throughout all the procedure. The ovaries were treated for four hours in permeabilization solution (PBS, 0.2% Polyvynal alcohol (PVA), 0.1% NaBH4-solution (Sigma) and 1.5% Triton X-100), than incubated for 24 hours in blocking solution (PBS, 0.1% Triton X-100, 0.15% Glycine pH7.4, 10% normal goat serum, 3% BSA, 0.2% sodium azide and 100 units/ml penicillin, 100 μg/ml streptomycin, and 0.25 μg/ml Fungizone). All the immunostaining and clearing was performed at room temperature (RT) with gentle rocking. Antibodies were diluted to appropriate concentration in the blocking solution. Primary antibodies (mouse anti-p63 (1:500, 4A4, Novus Biologicals); and rabbit anti-MVH (1:600, Abcam)) were incubated for four days. Afterwards, ovaries were washes with washing solution (PBS, 0.2% PVA and 0.15% triton X-100) for 10 hours than two times of two hours. Secondary antibodies (1:1000 Alexa Fluor® secondary antibodies) were incubated for three days in a vial protected from light. Ovaries were washes with washing solution for three times of 12 hours (if needed DAPI 50ng/ml was added to the first wash).

### Clearing, imaging, and oocyte quantification

Immunostained ovaries were cleared with modified, freshly-prepared ScaleS4(0) reagent (40% D-(-)-sorbitol (w/v), 10% glycerol, 4M urea, 20% dimethylsufoxide, pH8.1 (HAMA *et al.* 2015), gently mixed by inversion at 50°C for 30min and degassed prior to use). Solution was refreshed twice daily until tissue became transparent (usually two days). The insert was placed on top of a glass slide, and the membrane containing the cultured ovaries was carefully removed with a fine tip scalpel and placed on the slide. Slides were imaged on an upright laser scanning Zeiss LSM880 confocal/multiphoton microscope, using a 10X NA 0.45 water immersion objective. For proper image stitching the adjacent images (tiles) were overlapped by 20%. The z-steps were set for 5μm between optical sections. Images were reconstructed, visualized and analyzed using Fiji-ImageJ (SCHINDELIN *et al.* 2012).

Movies were made using the 3D project feature of Fiji-ImageJ (SCHINDELIN *et al.* 2012). Oocyte quantification was performed in flattened maximum intensity projections of the Z-stacks image-series, using the “analyze particle” feature of Fiji-ImageJ.

### Statistical analysis

Statistical analyses were done using JMP Pro12 software (SAS Inc., Cary, NC-USA, version 12.0.1). Fertility was analyzed using a mixed model with mother as random effect and ovary treatment as fixed effect. Least square (LS) means difference between litter sizes derived from treated vs non-treated ovaries was performed using the Student’s t test. LS mean differences between follicle counts from the different treatment groups were tested using Tukey honest significance different (HSD).

## REFERENCES

Antoni, L., N. Sodha, I. Collins and M. D. Garrett, 2007 CHK2 kinase: cancer susceptibility and cancer therapy - two sides of the same coin? Nat Rev Cancer 7:925–936.

Arienti, K. L., A. Brunmark, F. U. Axe, K. McClure, A. Lee et al., 2005 Checkpoint kinase inhibitors: SAR and radioprotective properties of a series of 2-arylbenzimidazoles. J Med Chem 48:1873–1885.

Bolcun-Filas, E., V. D. Rinaldi, M. E. White and J. C. Schimenti, 2014 Reversal of female infertility by Chk2 ablation reveals the oocyte DNA damage checkpoint pathway. Science 343:533–536.

Chehab, N. H., A. Malikzay, M. Appel and T. D. Halazonetis, 2000 Chk2/hCds1 functions as a DNA damage checkpoint in G(1) by stabilizing p53. Genes Dev 14:278–288.

Garrett, M. D., and I. Collins, 2011 Anticancer therapy with checkpoint inhibitors: what, where and when? Trends in pharmacological sciences 32:308–316.

Gonfloni, S., T. L. Di, S. Caldarola, S. M. Cannata, F. G. Klinger et al., 2009 Inhibition of the c-Abl-TAp63 pathway protects mouse oocytes from chemotherapy-induced death. Nature medicine 15:1179–1185.

Hama, H., H. Hioki, K. Namiki, T. Hoshida, H. Kurokawa et al., 2015 ScaleS: an optical clearing palette for biological imaging. Nat Neurosci 18:1518–1529.

Heyer, B. S., A. MacAuley, O. Behrendtsen and Z. Werb, 2000 Hypersensitivity to DNA damage leads to increased apoptosis during early mouse development. Genes Dev 14:2072–2084.

Hirao, A., Y. Y. Kong, S. Matsuoka, A. Wakeham, J. Ruland et al., 2000 DNA damage-induced activation of p53 by the checkpoint kinase Chk2. Science 287:1824–1827.

Kerr, J. B., K. J. Hutt, M. Cook, T. P. Speed, A. Strasser et al., 2012 Cisplatin-induced primordial follicle oocyte killing and loss of fertility are not prevented by imatinib. Nature medicine 18:1170–1172.

Kim, S. Y., S. K. Kim, J. R. Lee and T. K. Woodruff, 2016 Toward precision medicine for preserving fertility in cancer patients: existing and emerging fertility preservation options for women. J Gynecol Oncol 27:e22.

Lee, S. J., and J. Y. Wang, 2009 Exploiting the promiscuity of imatinib. J Biol 8:30.

Livera, G., B. Petre-Lazar, M. J. Guerquin, E. Trautmann, H. Coffigny et al., 2008 p63 null mutation protects mouse oocytes from radio-induced apoptosis. Reproduction 135:3–12.

Maiani, E., B. C. Di, F. G. Klinger, S. M. Cannata, S. Bernardini et al., 2012 Reply to: Cisplatin-induced primordial follicle oocyte killing and loss of fertility are not prevented by imatinib. Nature medicine 18:1172–1174.

Oktay, K., O. Oktem, A. Reh and L. Vahdat, 2006 Measuring the impact of chemotherapy on fertility in women with breast cancer. Journal of clinical oncology : official journal of the American Society of Clinical Oncology 24:4044–4046.

Pacheco, S., M. Marcet-Ortega, J. Lange, M. Jasin, S. Keeney et al., 2015 The ATM signaling cascade promotes recombination-dependent pachytene arrest in mouse spermatocytes. PLoS Genet 11:e1005017.

Redig, A. J., R. Brannigan, S. J. Stryker, T. K. Woodruff and J. S. Jeruss, 2011 Incorporating fertility preservation into the care of young oncology patients. Cancer 117:4–10.

Salama, M., and P. Mallmann, 2015 Emergency fertility preservation for female patients with cancer: clinical perspectives. Anticancer Res 35:3117–3127.

Schindelin, J., I. Arganda-Carreras, E. Frise, V. Kaynig, M. Longair et al., 2012 Fiji: an open-source platform for biological-image analysis. Nat Methods 9:676–682.

Suh, E. K., A. Yang, A. Kettenbach, C. Bamberger, A. H. Michaelis et al., 2006 p63 protects the female germ line during meiotic arrest. Nature 444:624–628.

Takai, H., K. Naka, Y. Okada, M. Watanabe, N. Harada et al., 2002 Chk2-deficient mice exhibit radioresistance and defective p53-mediated transcription. Embo J 21:5195– 5205.

Woodruff, T. K., 2007 The emergence of a new interdiscipline: oncofertility. Cancer Treat Res 138:3–11.

